# Myosin driven Actin Filament Sliding is Responsible for Endoplasmic Reticulum and Golgi Movement

**DOI:** 10.1101/347187

**Authors:** Joseph F McKenna, Stephen E D Webb, Verena Kriechbaumer, Chris Hawes

**Affiliations:** Plant Cell Biology, Dept. Biological and Medical Sciences, Oxford Brookes University, Oxford, OX3 0BP; Central Laser Facility, Research Complex at Harwell, Science and Technology Facilities Council, Rutherford Appleton Laboratories, Oxfordshire OX11 0QX; Biotechnology and Biological Sciences Research Council, Polaris House, Swindon, SN2 1UH

**Keywords:** Actin, Myosin, Endoplasmic Reticulum, Golgi, FRAP, Photoactivation

## Abstract

The plant secretory pathway is responsible for the production of the majority of proteins and carbohydrates consumed on the planet. The early secretory pathway is composed of Golgi bodies and the endoplasmic reticulum (ER) and is highly mobile in plants with rapid remodelling of the ER network. The dynamics of the ER and Golgi bodies is driven by the actin cytoskeleton and myosin motor proteins play a key role in this. However, exactly how myosin motor proteins drive remodelling in plants is currently a contentious issue. Here, using a combination of live cell microscopy and over-expression of non-functional myosins we demonstrate that myosin motor proteins drive actin filament sliding and subsequently the dynamics of the secretory pathway.

**Summary:** In plants, the actin cytoskeleton and myosins are fundamental for normal dynamics of the endomembrane system and cytoplasmic streaming. We demonstrate that this is in part due to myosin driven sliding of actin filaments within a bundle. This generates, at least in part, the motive force required for cell dynamics *in planta*.

## Introduction

Many organelles in plant cells display dynamic movements in the cytoplasm. At its simplest these can be divided into two classes. Firstly, classic cytoplasmic streaming, where movement is due to shear forces induced by actomyosin driving larger organelles [1]. This can be particularly apparent in *trans*-vacuolar strands of leaves. Secondly, more controlled movement of organelles over the actin cytoskeleton at the cortex of cells, especially in highly vacuolated tissues [2]. Even before the advent of live cell imaging based on fluorescent protein technology, it had been shown that in plants the endoplasmic reticulum (ER) network structure and Golgi body dynamics are somehow organised by the actin cytoskeleton [3,4]. By the combination of GFP expression and live cell imaging this organisation is exemplified by the movement of Golgi bodies and ER exit sites (ERES) with the ER network [2,5] and the various different movements of the ER, with four distinct forms having been identified [6–9]. Given that the ER and secretory pathway are the primary site of protein and carbohydrate production in plants, and hence the basis of global food supply, understanding the mechanism behind ER remodelling could have wide reaching implications. Both remodelling of the ER and Golgi movement are abolished by depolymerisation of actin, demonstrating the importance of the actin cytoskeleton in intracellular movement [2,7]. In addition, the cortical actin cytoskeleton supports organelle movement and is dynamic in its own right. Actin filaments and bundles continually remodel in the cytoplasm, and this can involve lateral-filament migration, sliding on actin bundles, filament severing and elongation [10–12].

Plants have two classes of myosins, VIII and XI. There are four members of the myosin VIII family, which most likely function as tensors at the cell surface rather than as motors [13]. In arabidopsis there are thirteen members of the myosin XI family most of which display some form of motor activity [14]. The fastest known myosin of *Chara corallina* reaches speeds of up to 50µms^-1^ in *in vitro* assays and is significantly faster than mammalian homologues [15,16]. In higher plant cells, myosin XI motility is up to 5µms^-1^, an order of magnitude lower than in algae but still 10-fold faster than the closest human homologue (myosin Va [17,18]). Mutant knock-out analysis of four members of the arabidopsis XI family (*xi-k, xi-1, xi2* and *xi-i*) demonstrate that these proteins are important for normal whole-organism and cellular growth as well as Golgi body dynamics [19]. They are also important for normal dynamics of the actin cytoskeleton *in planta*, with the knockout plants showing decreased turnover with a 2-fold reduced filament severing frequency [20]. Chimeric expression of a slow myosin XI-2, composed of the native myosin XI-2 with the motor domain replaced with *Homo sapiens* myosin Vb motor domain or a fast chimeric myosin protein which contains the *Chara corallina* myosin XI motor domain, results in smaller and larger cell size respectively [21]. This correlation between myosin motility and cell size demonstrates the importance of the actomyosin system in plants and its role in biomass production. Over-expression of truncated forms of the myosin XI family in tobacco leaf cells, show that a number of these will inhibit both Golgi and ER movement, presumably by complexing with the native myosin and rendering it non-functional. This could be in a similar manner to mammalian Myosin Va tail domain expression which turns Myosin Va into an inactive conformation [22].

A possible explanation for myosin generated movement is that myosin molecules link the organelles to the actin cytoskeleton and generate the necessary motive force to explain all the observed movements. However, despite a number of publications expressing fluorescently tagged myosin constructs of various different forms, there is no convincing evidence of any full length myosins decorating the ER surface or Golgi bodies [14,23]. It has however been suggested that myosin XI-K associates with endomembrane-derived vesicles in arabidopsis [24]. DIL domain (homologue of a yeast secretory vesicle binding domain of Myo2p) constructs from arabidopsis myosin XIs locate to various organelles and only one from myosin XI-G labels Golgi and ER [25]. Thus, apart from these few hints, it is still an open question as to how force is generated to induce motility in the two major organelles of the secretory pathway, the ER and Golgi bodies in conjunction with the actin cytoskeleton.

A possible mechanism to explain movements of the cortical ER network is that the motive force comes from motile Golgi bodies attached to the actin cytoskeleton [26]. Golgi bodies associated with ER networks are restricted to curved membrane surfaces in yeast [27] and higher plants [28]. It was demonstrated in tobacco leaf cells that if Golgi bodies were disrupted with Brefeldin A, resulting in the reabsorption of Golgi membrane back to the ER, ER motility still persisted [4]. It is however possible that a residual matrix of Golgi associated proteins and putative ER tethers remains after such experiments and could still remain associated with the actin cytoskeleton during such drug treatments [29,30].

Direct contacts between ER and actin have been reported [31,32]. Cao et al., (2016) [32] reported that the SNARE protein SYP73 may act as a linker between the ER, myosin and the actin cytoskeleton. Although an earlier report of the closely related ER SNARE SYP72 suggested that its function was to mark Golgi-ER import sites on the surface of the ER membrane, with no mention of any association with actin or actin binding proteins [33]. Likewise a protein of the NETWORKED family, NET 3B has been shown to bridge between the ER and actin bundles [34].

Another explanation for some of the observed ER/Golgi motility is that there is no direct connection between actin filaments, myosin and the organelles, even via linker or receptor proteins. Here we have used three different probes for plant actin, fABD2 [10,35] Lifeact [36,37] and a GFP tagged anti-actin alpaca nanobody [38] which, when expressed, self-immunolabels the actin cytoskeleton. We demonstrate that upon overexpression of a dominant negative myosin tail domain, Fluorescence Recovery After Photobleaching (FRAP) of all three actin marker labels is reduced. Furthermore, upon photoactivation of actin labelled lines, the signal migrates out of the activation area to adjacent actin filaments, and this is inhibited by overexpression of the dominant negative myosin tail domain. Our hypothesis is that ER remodelling is generated by myosin induced sliding of actin filaments over one another and that a non-motor link between actin filaments and ER membrane most likely transfers motive force to the ER membrane, perhaps via SNAREs or NET proteins such as SYP73 and NET3B [32,34].

## Results

### ER tubule elongation moves over existing actin bundles

The cortical ER in cotyledon and mature leaf epidermal cells shows various differing movements including polygon rearrangement, movement of the membrane surface and tubule outgrowths [7]. To test if the latter was mediated by actin filament polymerisation, we transiently co-expressed GFP-fABD2 with the ER marker ssRFP-HDEL and imaged with sub-diffraction-limited resolution using the Airyscan detector (Fig. 1; video 1). We found no evidence of ER tubules tracking actin filament polymerisation. However, growing tubules were routinely imaged moving along pre-existing actin bundles (Fig. 1 and video 1, white arrows).

**Figure 1.**
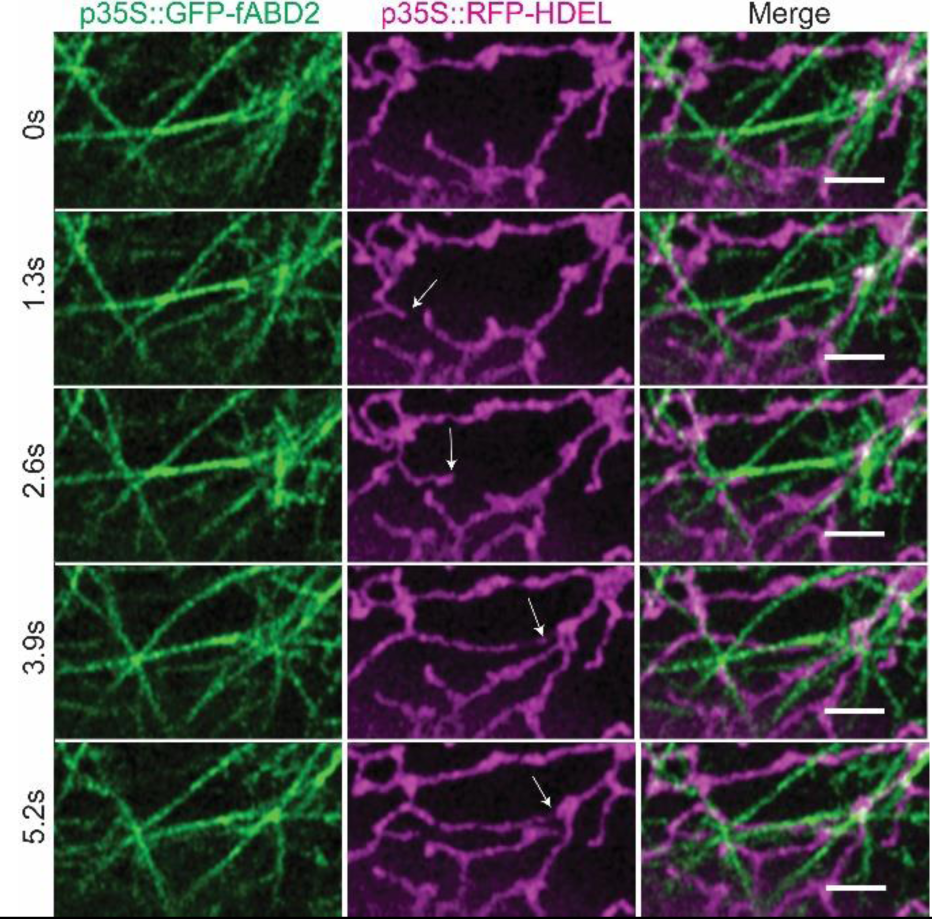
Airyscan imaging of an endoplasmic reticulum tubule elongating over existing actin filaments in *N. benthamiana*. N = 60 cells across 3 experimental repeats. Representative images shown. White arrows showing growing ER tubule. Scale bar = 2µm.

### FRAP recovery of labelled actin cytoskeleton demonstrates myosin dependence

We have previously shown that expression of non-functional myosin XI tail fragments can inhibit movement of Golgi bodies and the ER [26,39]. However, fragments from several of the 13 myosin XI isoforms, on expression in tobacco leaf cells, had no or negligible effects on endomembrane dynamics. We therefore combined our experiments with transient expression of myosin tail fragments in benthamiana leaf pavement cells, to assess the potential role of myosins in actin filament sliding within actin bundles. We decided not to use myosin inhibitor drugs such as BDM or ML-7 as their specificity in plants has been called into question [40,41]. Initially we used Total Internal Reflection Fluorescence (TIRF) microscopy to image actin filaments and bundles with high temporal and spatial resolution. In addition to the previously reported changes in actin cytoskeleton structure in myosin knockout mutant lines [1], actin filament dynamics were also impaired (Fig. 2A&B) when a dominant negative myosin tail domain (XI-K) was overexpressed (Fig.2A&B). Reductions in actin dynamics and more bundled networks have previously been reported for triple knockout mutants in arabidopsis [1,20]. Here we demonstrate that this is phenocopied in *N. benthamiana* with transient expression of a dominant negative myosin tail domain (Fig. 2). In order to determine in more detail how fluorescence recovery occurs, we performed FRAP experiments by TIRF microscopy. This recovery occurs along existing filaments (Fig. 2C), but not uniformly, as one would expect if recovery was due to new binding of GFP-fABD2 along the entire length of the filament. For actin labelled with fABD2-GFP and imaged using confocal microscopy, recovery after photobleaching was significantly impeded when myosin XI-K tail fragments were transiently expressed in the leaves (Fig. 3A-D; video 2). Control plateau and t1/2 values were 59.55 ± 17.7% (SD) and 4.6 ± 2s (SD) compared to XI-K values of 44.4 ± 17.6% (SD) and 6.1 ± 3.2s (SD) respectively (Fig. 3B-D) (p≤0.0001 for plateau, P≤0.001 for t1/2, ANOVA). On expression of a tail fragment of myosin XI-A, previously shown not to inhibit mitochondria movement [39] or ER remodelling [42], there was no significant effect on fluorescence recovery. Therefore, the observed effects with XI-K are not due to unspecific expression of tail domains, but only those which inhibit endomembrane dynamics (XI-K). This suggests that myosins contribute toward the recovery of labelled actin fluorescence by supporting inter-filament actin sliding. Photobleaching was also performed on the reporter GFP-Lifeact in combination with XI-K tail domain expression (Supplementary Fig. S1B-E) and the intensity plateau level was significantly reduced between the control and XI-K, and between XI-A and XI-K (Fig. S1). The t1/2 values for Lifeact photobleaching were not significantly different between the control and, XI-K or XI-A. To summarise, in two different actin marker lines, expression of the inhibitory tail domain of XI-K reduced actin dynamics.

**Figure 2.**
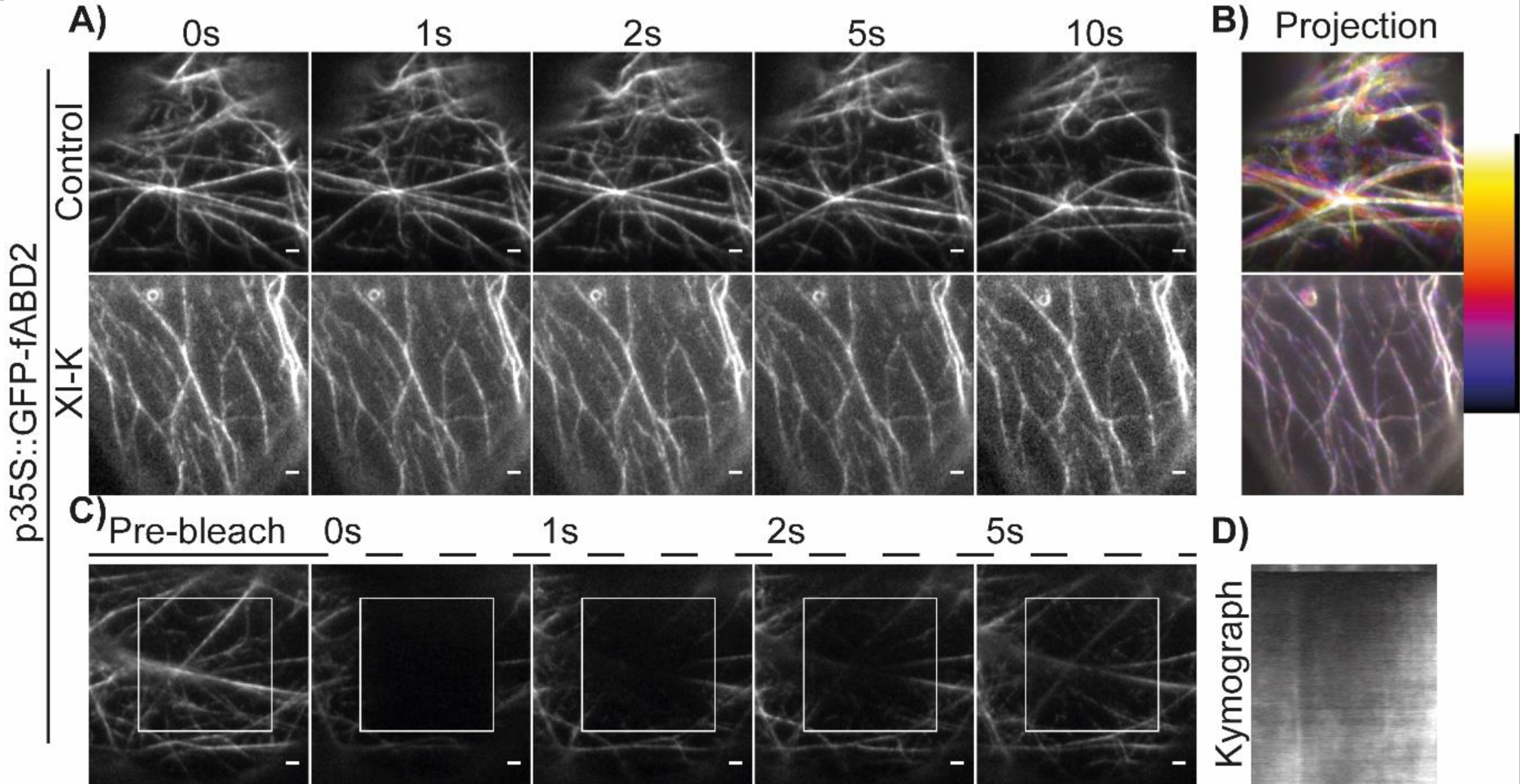
TIRF imaging and transient expression in *N. benthamiana* demonstrates dominant negative myosin XI-K tail domain perturbs actin dynamics *in planta*. A) TIRF Time course data from labelled GFP-fABD2 control and with p35S::RFP-XI-K. N=20 cells, representative images shown. B) Temporal colour coded projection of time-course data in A). C) Fluorescence recovery from the actin cytoskeleton labelled with GFP-fABD2 occurs along existing filaments. N=30 cells, representative images shown. D) Kymograph showing FRAP recovery along actin filament occurs from either side of the bleach region. Scale bar = 2µm.

**Figure 3.**
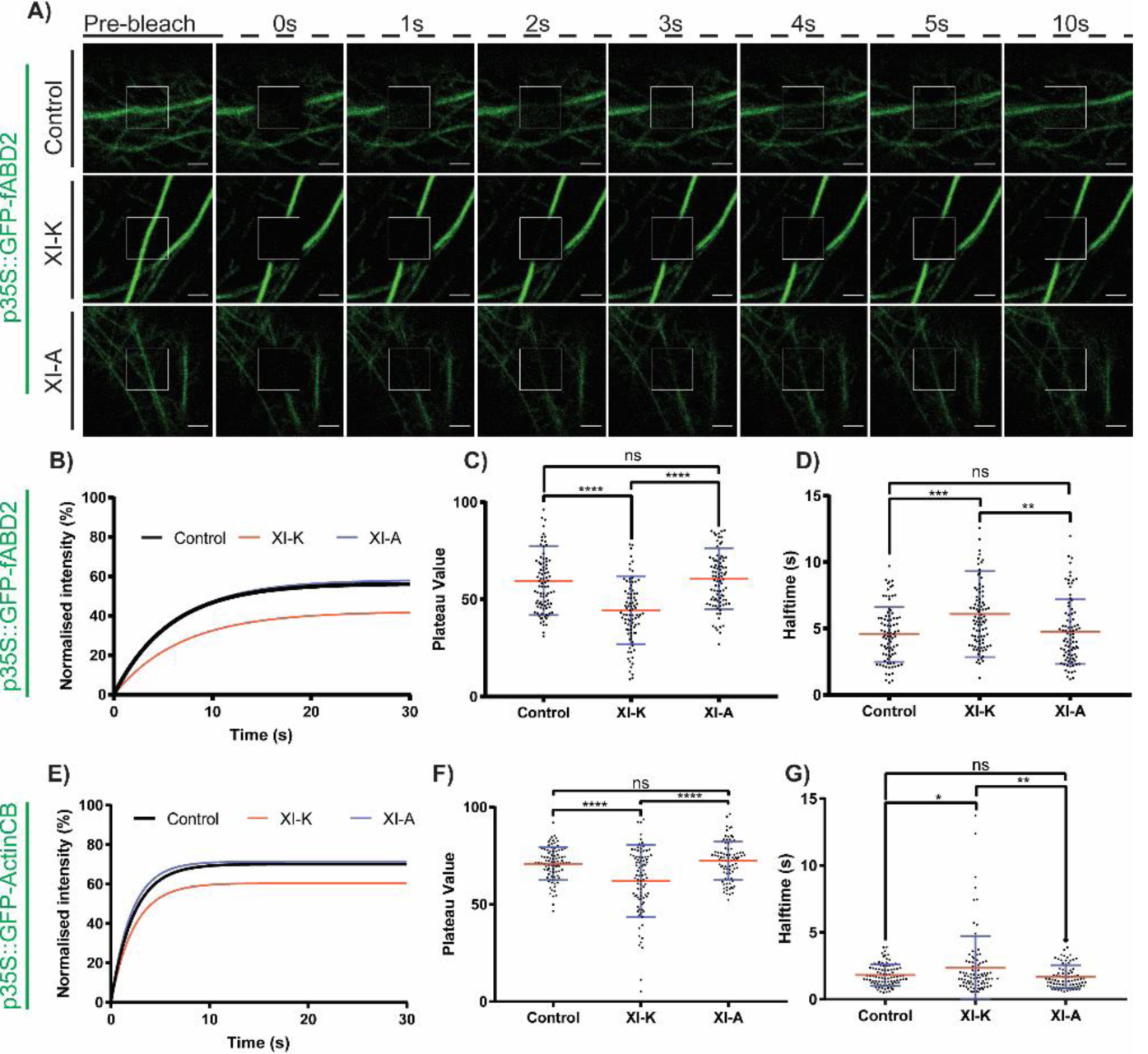
Inhibition of myosin via overexpression of a dominant negative tail domain reduces GFP-fABD2 labelled actin FRAP recovery in *N. benthamiana*. A) Time-course of confocal FRAP showing fluorescence recovery of GFP-fABD2 and co-expression with the dominant negative p35S::RFP-XI-K tail domain. The XI-A tail domain had no effect on ER remodelling or Golgi mobility and so was used as a negative control. B) Fluorescence recovery curves, C) plateau values, D) halftimes of fluorescence recovery for GFP-fABD2 FRAP from control, XI-K and XI-A treatment. E) Fluorescence recovery curves, F) plateau values, G) halftimes of fluorescence recovery for GFP-actinCB FRAP from control, XI-K and XI-A treatment. For boxplots, error bars (blue) denote standard deviation, mean value (red) is shown. ANOVA statistical analysis was performed. ns = p≥0.05, **=p≤0.01, ***=p≤0.001, ****=p≤0.0001. N=90 cells, across 3 experimental repeats. Scale bar = 2µm.

Similar experiments were carried out on the actin cytoskeleton in cotyledon epidermal cells self-immunolabelled by transient expression of an alpaca chromobody against actin [38] and tagged with GFP (Fig. 3E-G and S1A, GFP-actinCB). Inhibition of fluorescence recovery after bleaching in the presence of myosin XI-K was not as dramatic as with fABD2-GFP labelled actin but still significant (Fig. 3F&G). Plateau values were 70.9 ± 8.5% (SD) for the control and 62 ± 18.5% (SD) for XI-K, a statistically significant difference (p≤0.0001, ANOVA, Fig 1B and C). The t1/2 values were 1.8 ± 0.8s (SD) for control and 2.4 ± 2.3s (SD) for XI-K, also a statistically significant difference (p≤ 0.05, ANOVA, Fig. 1D).

### Actin labelled FRAP recovery is not due to actin polymerisation

As a further control we measured fluorescence recovery of actin bundles after treatment with jasplakinolide which has previously been used *in planta* to stabilise actin filaments [43,44]. It induces hyper actin polymerisation, resulting in the depletion of available G-actin in the cell and therefore inhibiting subsequent polymerisation [45,46]. Jasplakinolide treatment did not significantly alter the recovery period, plateau or t_1/2_ of GFP-fABD2 labelled actin filament bundles (Fig 4A-D). This indicates that recovery was not due to actin filament polymerisation into the bleached zone. This was the same for the GFP-actinCB after jasplakinolide treatment (Supplementary Fig. S2), with no statistically significant difference in plateau level or t_1/2_. However, there was a significant decrease in Golgi velocity on treatment with the drug (Fig. 4E), although at this stage we have no information as to whether this reflects on a secondary effect of the treatment.

**Figure 4.**
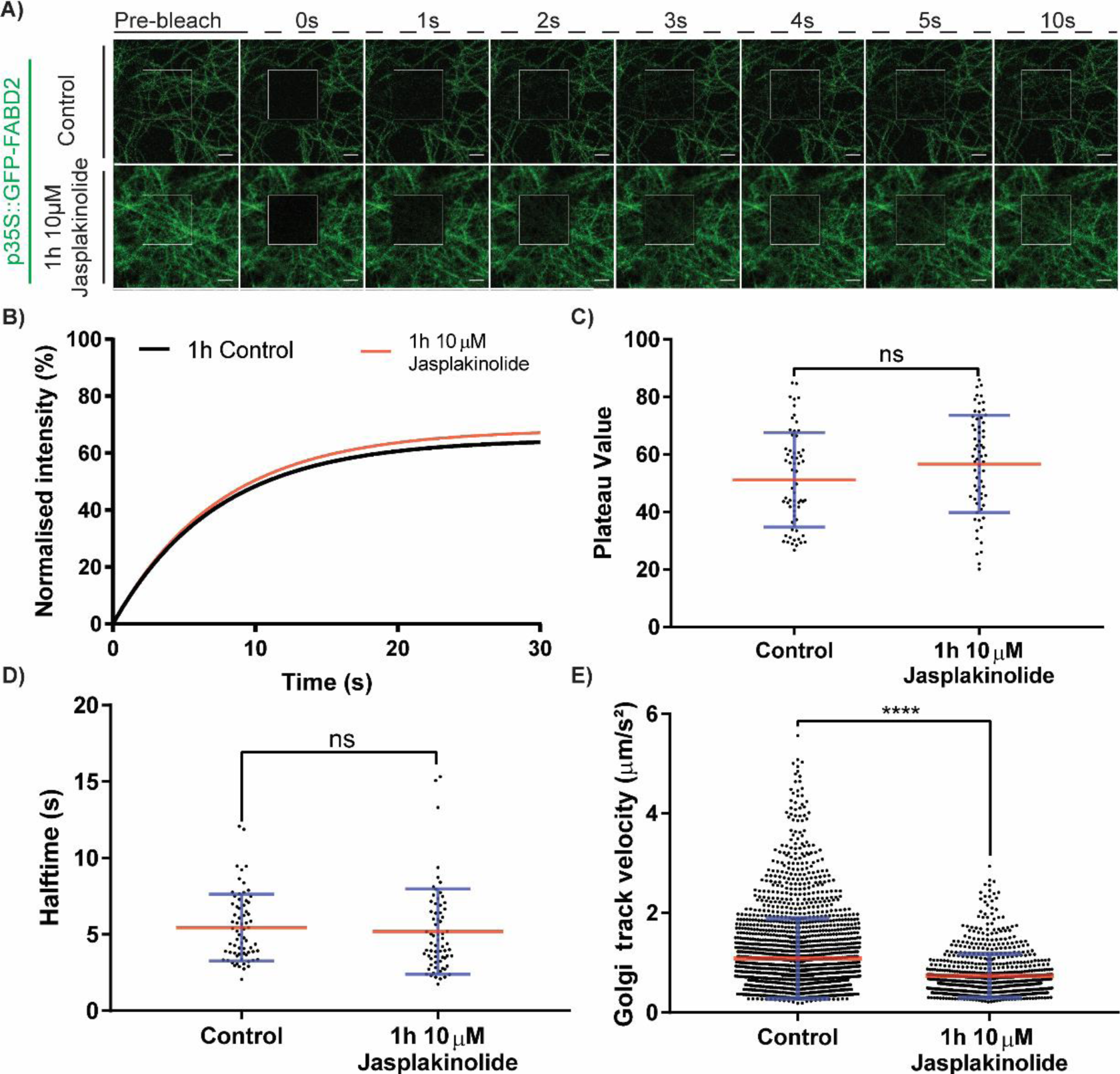
Treatment with Jasplakinolide and stabilisation of actin does not affect FRAP but does affect Golgi body mobility in *N. benthamiana*. A) Time-course of fluorescence recovery of GFP-fABD2 after Jasplakinolide treatment. N=≥63 cells across 3 experimental repeats per condition. B) Fluorescence recovery curves, C) plateau values and D) half-times of fluorescence recovery for control and 1h 10µM Jasplakinolide treated GFP-fABD2. E) Golgi track velocity of p35S::ST-RFP labelled Golgi bodies in 1h control and 10µM Jasplakinolide treated tissue. N=30 cells per condition. All experiments performed with transient infiltration in *N. benthamiana*. For boxplots, error bars (blue) denote standard deviation, mean value (red) is shown. Student T-test statistical analysis performed. ns = p≥0.05. ****=p≤0.0001. Scale bar = 2µm.

### Photoactivation demonstrates myosin dependant sliding of actin filaments

Alongside the photobleaching experiments, we confirmed our results by transiently expressing fABD2 or Lifeact linked to mCherry and photoactivatable-GFP (paGFP). In this way, it was possible to quantify the dispersal of the activated GFP along actin filaments (Fig.5A-E, arrows; video 3). On activation of the mCherry-paGFP constructs, the green fluorescence signal rapidly dispersed laterally over the mCherry labelled filaments (Fig. 5A; video 3). Measuring the intensity increase of an adjacent ROI a set distance from the activated region, an increase in GFP fluorescence occurred above the initial activation level after timepoint 0s (Fig. 5B). Both the plateau value and t_1/2_ of the activated ROI are significantly higher when expressed with XI-K than in the control or XI-A (control plateau: 5.9 ± 5.7% (SD), XI-K plateau 11.3 ± 13.4% (SD), Fig. 5D, control t^1/2^ 3.1 ± 1.3s (SD), XI-K t^1/2^ 5.1 ± 3.5s (SD), Fig. 5E). XI-A photoactivation was similar to the control, demonstrating the specificity of XI-K expression to filament recovery. In addition, mCherry-paGFP-Lifeact activation showed a slower t_1/2_ when expressed with XI-K than in the control or XI-A condition (Supplementary Fig. S3). Therefore, activated paGFP labelled actin moved more slowly out of the ROI when XI-K was expressed. This further demonstrated that sliding of actin filaments within actin bundles is regulated by myosins.

**Figure 5.**
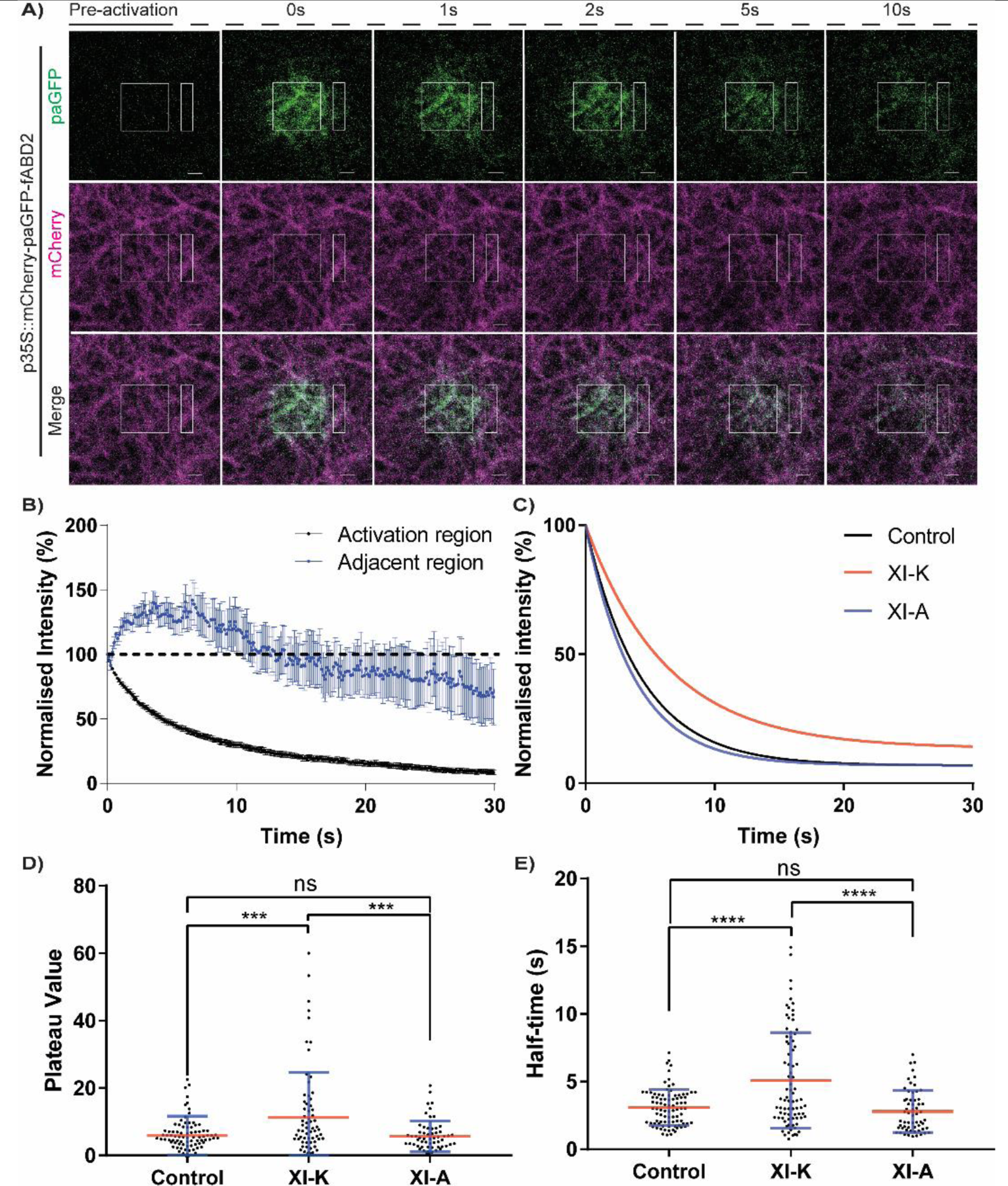
Photoactivation of dual labelled fABD2 lines demonstrates filament sliding is driven by myosins in *N. benthamiana*. A) Time-course of activation in control data. Activated region (central square) and adjacent non-activated region (white rectangle) are shown. mCherry (magenta) and paGFP (green) channels shown. B) Normalised intensity curve for activation region (black) and adjacent region (blue) (error bars = SE). C) One phase-decay plot for control, XI-K and XI-A expressing cells labelled with the dual marker. D) Plateau value from one-phase decay plot (C). E) Half-time of photoactivated actin marker. For boxplots, error bars (blue) denote standard deviation, mean value (red) is shown. ANOVA statistical analysis was performed. ns = p≥0.05, ***=p≤0.001, ****=p≤0.0001. N=≥60 across 3 experimental repeats. Scale bar = 2µm.

## Discussion

### Myosins are responsible for actin filament sliding

There are a number of ways in which the actin cytoskeleton can support movement within eukaryotic cells. These include the interaction between myosin motors, organelles and actin filaments; the rapid polymerisation of actin filaments and the myosin-driven sliding of actin filaments over each other within actin bundles. Plant myosins are now well documented and have long been known to support cytoplasmic streaming in a wide range of cells [47]. However, regarding the secretory pathway, the role of actin in the dramatic remodelling of the cortical ER network [1,3,7,42] and the movement of individual Golgi stacks [2,48–50] has been the subject of a number of reports. It has also been noted that Golgi bodies move in concert with the moving bounding membrane of the ER [6,51]. Although it is clear that members of the myosin XI family are involved in such movements [14,39,42,52], there is no convincing evidence for direct endomembrane organelle-myosin-actin interactions, with the exception that myosin XI-K constructs potentially labelling some post-Golgi compartments in roots. It was also suggested XI-K labelled ER derived vesicles in leaves with some ER labelling [24]. Furthermore, in a quadruple myosin mutant knockout line, the actin cytoskeleton structure is altered in addition to organelle dynamics [19]. Additionally, fluorescently tagged, myosin XI-K labels the actin cytoskeleton preferentially to post-Golgi compartments, as determined by co-localisation [19].

Here we demonstrate that new ER tubule formation occurs predominantly along existing actin bundles (Fig. 1). This demonstrates that either myosins moving along existing actin filaments or inter-filament sliding of actin dragging newly forming ER tubules occurs. We then used photobleaching and photoactivation to determine if myosin driven actin filament sliding occurs. The interpretation of fluorescence recovery data from fluorescent protein-tagged actin networks in plants can be fraught with problems. Both the commonly used fABD2 and Lifeact constructs bind to actin filaments and are subject to on and off turnover on the filaments themselves [53]. Thus, data from bleaching experiments can either be interpreted as a measurement of turnover of the actin binding fragments or movement/recovery of the actin filaments/bundles themselves. To mitigate this problem, we also utilised actin labelling by the expression of a cameloid actin nanobody spliced to GFP. This results in self-immunolabelling *in vivo* of the actin network [38]. Being an antibody fragment, its turnover rate on the actin filaments is very low. The results obtained were the same as those with the classic actin markers Lifeact and fABD2, demonstrating that myosin perturbation affects the recovery of fluorescence of the labelled actin cytoskeleton. This further supports the hypothesis that myosin drives filament sliding within actin bundles. Over-expression of a non-inhibiting XI-A myosin tail fragment had no effect on FRAP recovery of the actin labels used, therefore the results obtained are specific to ER and Golgi regulating myosins.

Jasplakinolide stabilises the actin cytoskeleton and depletes the pool of G-actin thereby not allowing subsequent polymerisation. After jasplakinolide treatment, we did not see an effect on photobleaching recovery of fluorescently labelled actin (Fig. 4), which is further proof that the recovery we observed is due to myosin driven filament sliding and not polymerisation. Furthermore, while there is a reduction in Golgi body velocity after jasplakinolide treatment, they are still moving and the actin network structure is perturbed. The decrease in movement could be due to network structure changes. As cytoskeleton FRAP recovery still occurs and Golgi bodies are still mobile this implies that myosin driven filament sliding contributes to their mobility and hence ER remodelling.

In parallel, we also employed photoactivatable constructs to label the cortical actin cytoskeleton, which report on the movement of activated fluorescence *and* the actin network simultaneously, not simply the turnover of constructs on the filaments. This photoactivation strategy clearly demonstrates movement of fluorescence (and hence bound actin) out of the activated regions into adjacent ones along existing actin filaments. This clearly demonstrates actin filament sliding. Furthermore, upon co-expression with a dominant negative myosin-tail domain, the loss of paGFP fluorescence out of the activation region is reduced, demonstrating that the filament sliding observed is, at least in part, myosin dependant.

## Summary

Utilising the photobleaching and activation strategy our results demonstrate that the cortical actin filaments within bundles are sliding over each other powered by one or more myosins. To support movement of the ER and Golgi bodies attached to it, it would be necessary to anchor the ER membrane to underlying actin filament bundles. Several recent reports have suggested that SNARE proteins (SYP 73, [32]) and members of the NETWORKED family, NET3b over the ER [34] and NET3c at ER-plasma membrane contact sites [54], may perform this role. Myosin regulation of actin network structure has been reported previously, with the mammalian myoX motor function being critical for actin reorganisation at leading edges, resulting in filopodia formation [55]. In addition, the mammalian myosin1c stabilises ER sheets via regulation of actin filament array organisation [56]. Furthermore, it has been demonstrated *in planta* that myosins are responsible for generating the force required for buckling and straightening of both individual filaments and bundles [20]. Elegant work using optical tweezers has also demonstrated a role for myosin in actin entry into generated cytoplasmic protrusions [57]. Both of these *in planta* observations hypothesized that myosin facilitated sliding of filaments could account for this, which our work demonstrates.

We propose a new model that both ER and Golgi movement are, at least in part, a result of myosin driven sliding of actin filaments within actin bundles that underlie and are anchored to the ER (Fig. 6). This model can account for differences in speeds of ER and Golgi movement. Myosin motors and actin filaments can act independently or synergistically with each other to induce a range of different speeds of sliding, resulting in differential movement of the filaments attached to the ER. Indeed this model could also explain the differences in cell size observed when expressing fast and slow chimeric myosins [21] as these would result in respectively increased and reduced filament sliding and cytoplasmic streaming, hence larger and smaller cells. It can also explain the perturbed actin cytoskeleton observed in triple and quadruple mutant knockout lines. Furthermore, if there are different polarities of actin filaments within a bundle then directionality of ER membrane and associated organelle movement can be controlled in this manner.

**Figure 6.**
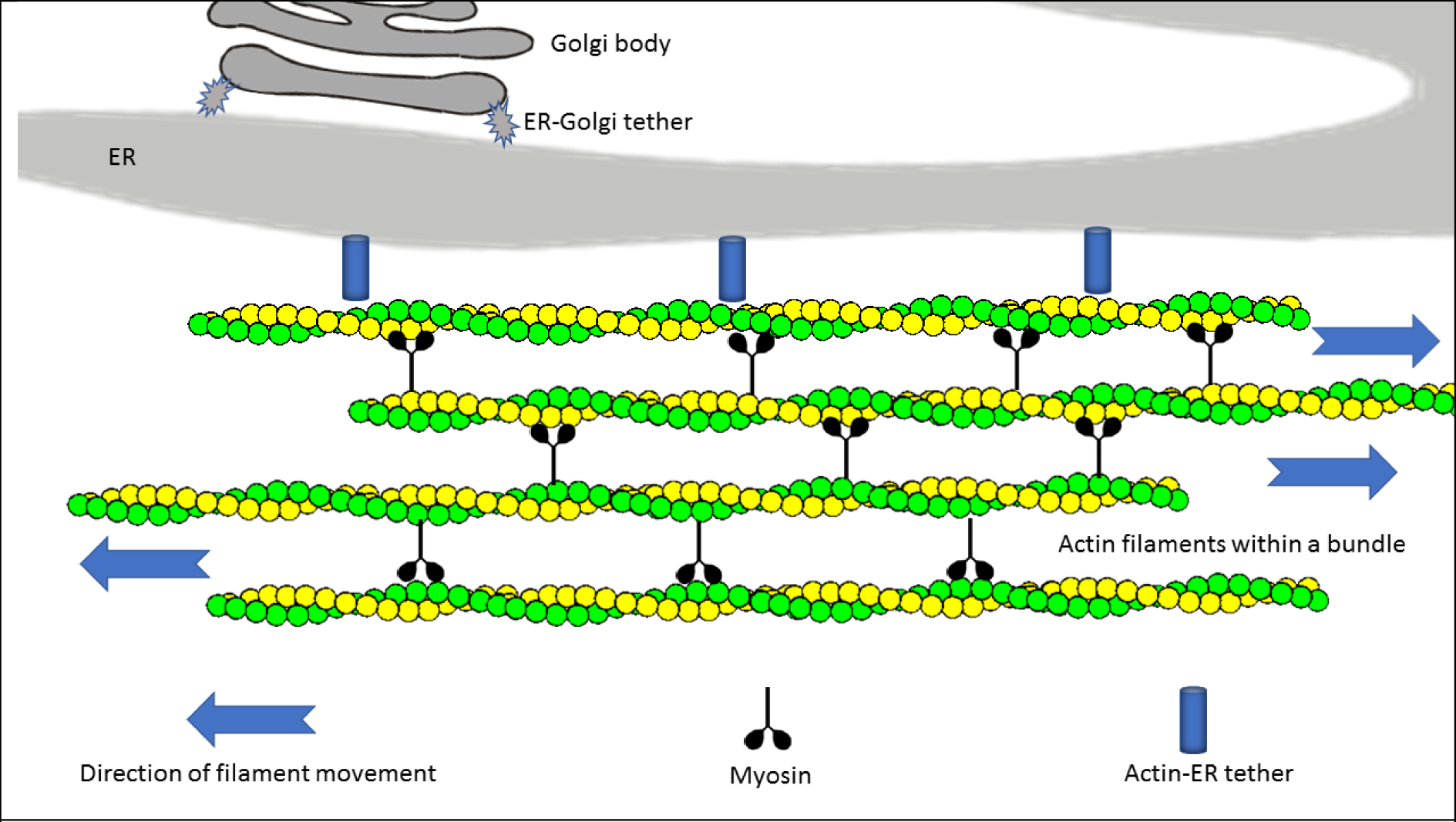
A model showing how myosin driven actin sliding with the combination of tethering proteins can potentially drive ER and Golgi mobility. Myosins are shown linking actin filaments within a bundle and are responsible for filament sliding. Potential linker proteins (such as SYP73) are shown tethering the ER to the actin cytoskeleton [32]. The Golgi bodies are tethered to the ER [50] and hence the physical force generated by actin sliding accounts for ER remodelling and Golgi body movement.

## Acknowledgements

We thank Dr Mark Fricker for helpful discussions and Dr Frances Tolmie for useful comments on the manuscript. We thank the Oxford Brookes Bioimaging unit for financial support. Access to the TIRF iLas system was provided under an STFC facility access grant to JM and SW (16230034).

## Author contributions

JM and CH conceived the experiments and wrote the manuscript. CH secured funding. JM and SEDW performed the experiments. VK provided genetic resources. All authors reviewed and edited the manuscript.

---

**Supplemental Figure 1.**
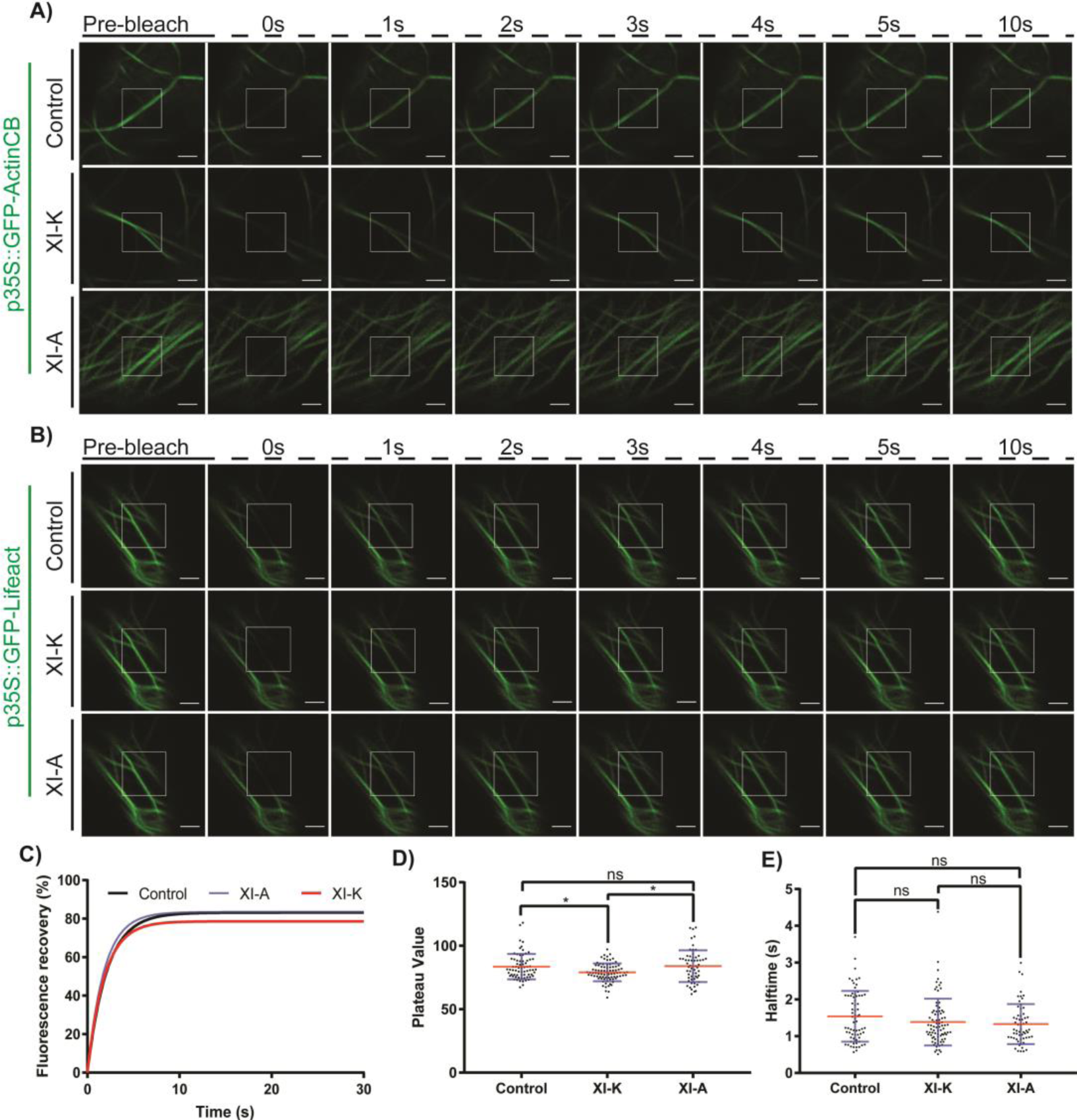
GFP-ActinCB and GFP-Lifeact FRAP with control, XI-K and XI-A expression transiently expressed in *N. benthamiana*. A) FRAP time course of GFP-actinCB labelled actin bundles in control, XI-K and XI-A conditions. FRAP quantification data in Fig. 3. B) FRAP time course of GFP-Lifeact labelled actin bundles in control, XI-K and XI-A conditions. B) Fluorescence recovery curves, C) plateau values and D) t^1/2^ of fluorescence recovery of actin bundles labelled with GFP-Lifeact shown in (A). For boxplots, error bars (blue) denote standard deviation, mean value (red) is shown. ns = p≥0.05, *=p≤0.05, ANOVA. N=≥59 cells, across 3 biological repeats per condition. Scale bar=2µm.

**Supplemental Figure 2.**
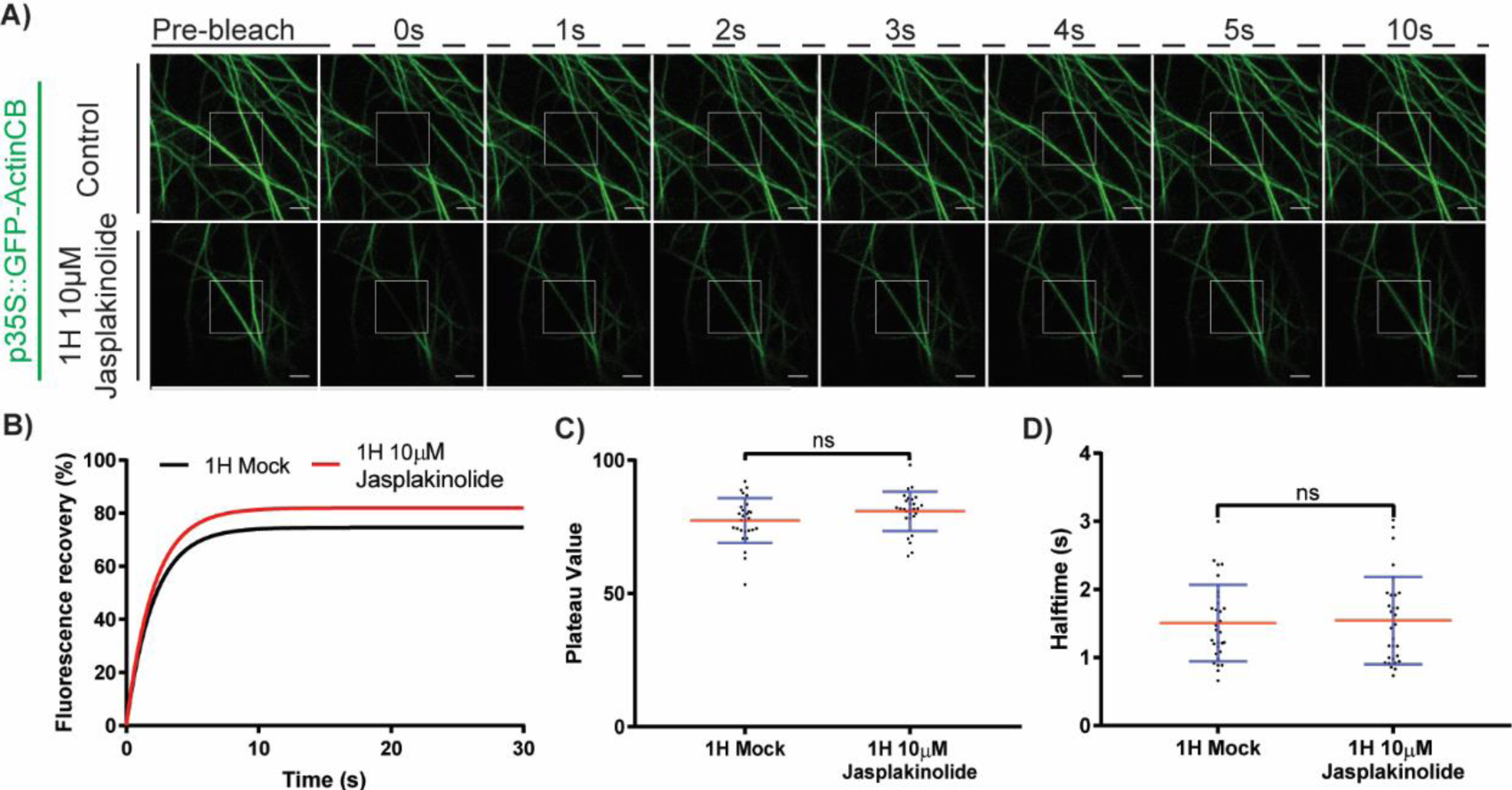
GFP-ActinCB FRAP recovery after treatment with jasplakinolide in transiently expressing *N. benthamiana*. A) FRAP time course of GFP-ActinCB labelled actin in 1h control and jasplakinolide treatments. B) Fluorescence recovery curves, C) plateau values and D) t^1/2^ of fluorescence recovery of actin shown in (A). For boxplots, error bars (blue) denote standard deviation, mean value (red) is shown. ns = p≥0.05, Students T-test. N=28 cells per condition.

**Supplemental Figure 3.**
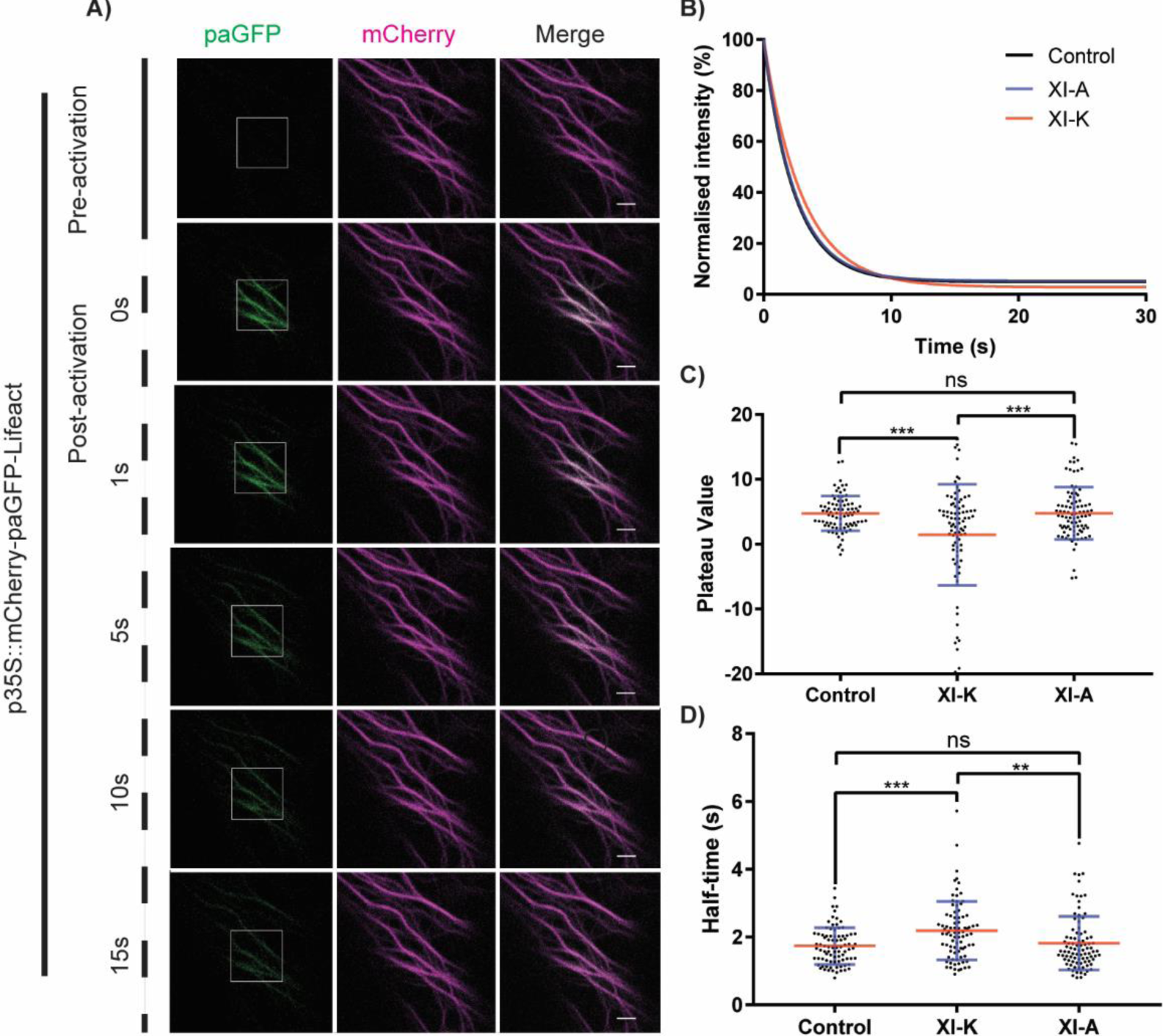
Photoactivation of mCherry-paGFP-Lifeact in control, XI-K and XI-A in transiently expressing *N. benthamiana* cells. A) Photoactivation of mCherry-paGFP-Lifeact labelled actin shows activated GFP sliding over actin filaments. B) Fluorescence decay curves, C) plateau values and D) t^1/2^ of fluorescence decay of control, XI-K and XI-A activated paGFP in (A). For boxplots, error bars (blue) denote standard deviation, mean value (red) is shown. ns = p≥0.05, ** = p≤0.01, *** = p≤0.001, ANOVA multiple comparison test. N=≥68 cells, across 3 experimental repeats per condition.

## Supplementary table 1

### Primers used in this study

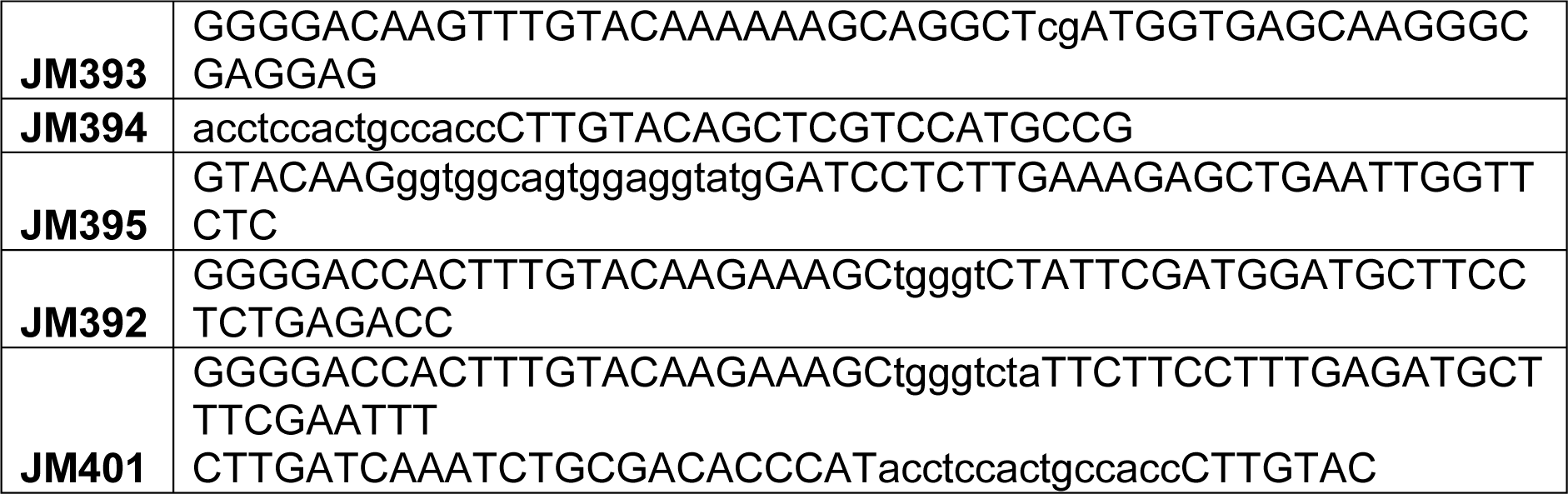

**Movie 1:** Time lapse series of ER tubules (labelled with p35S::RFP-HDEL) elongating over existing actin bundles (labelled with p35S::GFP-fABD2) in *N. benthamiana* transiently expressing leaf epidermal cells. Images same as in Fig. 2. White arrows show tip of elongating tubule. Scale bar denotes 2µm.

**Movie 2:** Time lapse series showing Flourescence recovery after photobleaching (FRAP) of GFP-fABD2 labelled actin expressed solo and with the myosin tail domains of XI-A or XI-K. Construct expression is transient in *N. benthamiana* leaf epidermal cells. White box indicates bleach region. Images same as in Fig. 4. Scale bar denotes 2µm.

**Movie 3:** Time lapse series showing photoactivation of paGFP of mCherry-paGFP-fABD2 labelled actin cytoskeleton bundles, demonstrating filament sliding. Construct expression is transient in *N. benthamiana* leaf epidermal cells. White box indicates activation region. Images same as in Fig. 7. Scale bar denotes 2µm.

## STAR* Methods

STAR\* Methods
| REAGENT or RESOURCE | SOURCE | IDENTIFIER |
| --- | --- | --- |
| **Bacterial and Virus Strains** |  |  |
| *E. Coli* strain DH5α | Widely distributed | N/A |
| *A. tumefaciens* strain GV3101 | Widely distributed | N/A |
| **Chemicals, Peptides, and Recombinant Proteins** |  |  |
| Q5® High-Fidelity DNA Polymerase | NEB | Cat# M0491S |
| NEBUILDER® HiFi DNA Assembly Master Mix | NEB | Cat# E2621S |
| MES hydrate | Sigma | Cat# M8250-100G |
| triSodium orthophosphate (Na<sub>3</sub>PO<sub>4</sub>) | BDH | Cat# 1777680 |
| Acetosyringone (4'-Hydroxy-3',5'-dimethoxyacetophenone) | Sigma | Cat# D134406 |
| D-Glucose | Fisher | Cat# G/0500/53 |
| Jasplakinolide | Cambridge bioscience | Cat# CAY11705-50ug |
| Spectinomycin | Melfords | Cat# S23000-25 |
| Rifampicin | Melfords | Cat# R0146 |
| Gentamicin | ThermoFisher | Cat# 15710-049 |
| BP Clonase | Invitrogen | Cat# 56480 |
| LR Clonase | Invitrogen | Cat# 56485 |
| **Critical Commercial Assays** |  |  |
| Nucleospin®-plasmid | Macherey-Nagel | Cat# 740588.250 |
| Nucleospin®- Gel and PCR cleanup | Macherey-Nagel | Cat# 740609.250 |
| **Experimental Models: Organisms/Strains** |  |  |
| *N. benthamiana* | Widely available | N/A |
| *A. tumefaciens* strain GV3101 p35S::RFP-XI-A | [39] | N/A |
| *A. tumefaciens* strain GV3101 p35S::RFP-XI-K | [39] | N/A |
| *A. tumefaciens* strain GV3101 p35S::GFP-fABD2 | [10] | N/A |
| *A. tumefaciens* strain GV3101 p35S::GFP-Lifeact | [35,37] | N/A |
| *A. tumefaciens* strain GV3101 p35S::GFP-ActinCB | This study | N/A |
| *A. tumefaciens* strain GV3101 p35S::mCherry-paGFP-fABD2 | This study | N/A |
| *A. tumefaciens* strain GV3101 p35S::mCherry-paGFP-Lifeact | This study | N/A |
| **Oligonucleotides** |  |  |
| A list of oligonucleotides is given in Table S1 |  |  |
| **Recombinant DNA** |  |  |
| pDONR221 | Invitrogen | Cat# 35-1687 |
| pB7WGF2 | VIB, Gent | N/A |
| pB7WGC2-mCherry | Modified from VIB, Gent, provided by John Runions, Oxford Brookes university. |  |
| p35S::GFP-ActinCB | This study | Modified from Chromotek ActinCB® https://www.chromotek.com/products/chromobodies/actin-chromobodyr/actin-chromobodyr-plasmid/ |
| p35S::mCherry-paGFP-fABD2 | This study | N/A |
| p35S::mCherry-paGFP-Lifeact | This study | N/A |
| **Software and Algorithms** |  |  |
| Fiji | https://fiji.sc/ | v1.52h |
| Zeiss Zen Black | https://www.zeiss.com/microscopy/int/products/microscope-software/zen-lite.html | v2.1 SP3 14.0.0.0. |
| Zeiss Zen Blue | https://www.zeiss.com/microscopy/int/products/microscope-software/zen-lite.html | v2.3 |
| Graphpad | https://www.graphpad.com/scientific-software/prism/ | v7.04 |
| Adobe illustrator CC | https://www.adobe.com/uk/products/illustrator.html | v22.0.1 |

## Experimental model and subject details

### Plant lines used and chemical treatments

N. *benthamiana* transient transformation was performed as described in Sparkes et al., [58]. All fluorophore-labelled marker lines were infiltrated with an *Agrobacterium* optical density at 600nm (OD) of 0.05. p35S::RFP-XI-A and p35S::RFP-XI-K [39] were transformed with an OD of 0.01. Imaging was performed at 3 days post infiltration (dpi). A Jasplakinolide stock solution (10mM) in DMSO was diluted to a working concentration of 10 µM in dH2O. Plant samples were incubated for 1h in Jasplakinolide or control (same concentration of DMSO) solutions.

## Method details

### Cloning constructs

#### GFP-ActinCB

The Actin-Chromobody^®^ plasmid containing the alpaca actin-antibody gene was obtained from Chromo-Tek (Martinsried, Germany [38]). Primers were ordered from Eurofins MWG Operon (Ebersberg, Germany). Q5 highfidelity DNA polymerase (New England Biolabs, Herts, UK) was used for the polymerase chain reaction (PCR) reaction. The ActinCb-PCR product was cloned into the binary vector pB7WGF2 providing an N-terminal GFP-tag using Gateway® technology (Invitrogen life sciences). For transient expression the construct was transformed into the *A. tumefacians* GV3101 strain under selection for spectinomycin, gentamycin and rifampicin.

#### mCherry-paGFP

Primer sequences are in supplemental table 1. For p35S::mCherry-paGFP-fABD2 primers were designed to amplify paGFP from the p35S::CXN-paGFP vector (Runions et al., 2006 [6]). The N-terminal FRD primer (JM393) was flanked with Gateway attB1 site and the C terminal REV primer (JM394) with a GGSGG amino acid linker overhang. fABD2 was then amplified by PCR using arabidopsis cDNA from five day old seedlings. The N-terminal FRD primer (JM395) consisted of a 22bp overhang composed of the GGSGG linker and last 7nt from paGFP. The C-terminal REV primer (JM392) contained a Gateway attB2 site. These DNA fragments were then fused together using the NEB HiFi Gibson assembly protocol. In order to generate p35S::mCherry-paGFP-Lifeact, paGFP fused to Lifeact was amplified from paGFP using an N-terminal primer (JM393), flanked with a Gateway attB1 site and a C-terminal REV primer (JM401) with an overhang composing a GGSGG linker, Lifeact and an attB2 site. Both constructs were then cloned into Gateway pDONR221 vector using BP clonase and subsequently a 35S promoter driven gateway compatible mCherry N-terminal vector using LR clonase to give p35S::mCherry-paGFP-fABD2 and p35S::mCherry-paGFP-Lifeact. These were then transformed into *A. tumefacians* GV3101 and selected for with Spectinomycin (50µg/ml), Gentamycin 15 µg/ml and Rifampicin 25µg/ml.

### Live cell microscopy

#### Confocal

Imaging was performed on Zeiss 880 or 800 confocal microscopes both equipped with Airyscan detectors and 100X 1.46NA lenses. Samples were mounted on #1.5 coverslips with dH2O. Airyscan imaging was performed on the Zeiss 880 using a 5X digital zoom and a 500-550BP and 565LP dual emission filter. An additional 620SP filter was used to block chlorophyll autofluorescence. For GFP and RFP/mCherry imaging, the 488 nm and 561 nm lasers respectively were used for excitation and the frame integration time was 0.13s. For GFP/RFP imaging, line switching was used (halving the frame rate). A minimum timeseries of 240 frames (≥30s) was collected for each FRAP and activation experiment. The FRAP experimental sequence was five pre-bleach image scans followed by 10 bleaching scans with the 488 nm laser at 100% in a square region (160×160 pixels) and then confocal imaging as described above. For photoactivation experiments, the 405 nm laser was used at 50% power for 10 iterations in a similarly square region to FRAP prior to imaging.

#### TIRF

TIRF imaging was performed on a Nikon Ti-E microscope equipped with an iLas2 TIRF FRAP system (Roper), Triline laserbank (Cairn), HQ525/50m emission filter (Chroma) and sCMOS detector (Prime 95B, Photometrics). A 100X 1.46NA lens was used and data was collected using MetaMorph. Excitation and bleaching were performed with a 488nm laser, with FRAP experiments involving 10 iterations of the 488nm laser at full power in the selected region of interest (ROI).

### Image analysis

Airyscan processing was performed in Zen Black version 2.1 SP3 14.0.0.0. ROI intensity data was extracted using Zen blue (v2.3). Image editing, kymographs and temporal colour coded projections was performed in FIJI (Image J version 1.51u). Golgi tracking was performed using Trackmate (v3.6.0) [59]. The RFP channel was segmented using a LoG detector with an estimated puncta diameter of 1µm, threshold of 10, a medium filter and sub-pixel localisation. All Golgi bodies tracked for fewer than 5 frames were discarded. FRAP analysis was performed as described in [60] with data being normalised and then fit to a non-linear regression one phase association curve. Photoactivation intensity data was normalised in the same way, however a non-linear regression one phase decay curve was fitted. For FRAP and photoactivation data, as well as recovery / decay curves, t_1/2_ and fluorescence plateau values were calculated. In order to demonstrate actin sliding, the intensities in the activation region and an adjacent region a set distance apart were analysed and normalised to T0 = 100% fluorescence intensity.

## Quantification and statistical analysis

Quantification of images was performed using either FIJI or Zen Blue. Data was collated in Microscoft excel. For graph generation and statistical analysis Graphpad was used. For reasons of clarity the statistical test performed (either ANOVA or t-test) and number of N for each experiment is listed in the corresponding figure legend. Significant difference is defined as: ns ≥ 0.05; * ≤ 0.05; ** ≤ 0.01; *** ≤ 0.001; **** ≤ 0.0001 and is indicated by asterisks above each box-plot. For box plots, blue error bars indicate the standard deviation (SD) and the red line represents mean value.

